# A principal component approach to improve association testing with polygenic risk scores

**DOI:** 10.1101/847020

**Authors:** Brandon J. Coombes, Joanna M. Biernacka

## Abstract

Polygenic risk scores (PRSs) have become an increasingly popular approach for demonstrating polygenic influences on complex traits and for establishing common polygenic signals between different traits. PRSs are typically constructed using pruning and thresholding (P+T), but the best choice of parameters is uncertain; thus multiple settings are used and the best is chosen. This optimization can lead to inflated type I error. To correct this, permutation procedures can be used but they can be computationally intensive. Alternatively, a single parameter setting can be chosen *a priori* for the PRS, but choosing suboptimal settings result in loss of power. We propose computing PRSs under a range of parameter settings, performing principal component analysis (PCA) on the resulting set of PRSs, and using the first PRS-PC in association tests. The first PC reweights the variants included in the PRS with new weights to achieve maximum variation over all PRS settings used. Using simulations, we compare the performance of the proposed PRS-PCA approach with a permutation test and *a* priori selection of p-value threshold. We then apply the approach to the Mayo Clinic Bipolar Disorder Biobank study to test for PRS association with psychosis using a variety of PRSs constructed from summary statistics from the largest studies of psychiatric disorders and related traits. The PRS-PCA approach is simple to implement, outperforms the other strategies in most scenarios, and provides an unbiased estimate of prediction performance. We therefore recommend it to be used PRS association studies where multiple phenotypes and/or PRSs are being investigated.

## Introduction

Polygenic risk scores (PRSs) have become an increasingly popular tool in genetics research. PRSs leverage summary statistics from previous genome-wide association studies (GWASs) to predict risk for individuals in a new population. If the individuals’ predicted risk is associated with their phenotype, this approach provides evidence of polygenetic effect even when no genome-wide significant variants exist. When a PRS for one trait is associated with another trait, this approach can be used to establish common polygenic signals between two different traits^1^.

A PRS is a weighted sum of an individual’s alleles, where allele weights are estimated based on their effects in a GWAS in a different sample^2^. A simple summation across single nucleotide polymorphisms (SNPs) while ignoring the linkage disequilibrium (LD) among them would not be appropriate because trait-associated regions with high LD would be over-weighted. There are several approaches that account for LD in PRS construction. The most common approach, the so-called “pruning-and-thresholding” (P+T) method, constructs the PRS by first removing SNPs in high LD to obtain a set of roughly independent SNPs (pruning) and then including only SNPs that have a p-value below a certain value (thresholding)^2,3^. Other methods use penalization to shrink most of the SNP effects to zero^4^ or use a Bayesian prior that incorporates the LD structure to place downward bias on all of the SNP effects^5,6^.

Regardless of the method, construction of a PRS requires specification of tuning parameters, such as the pruning and thresholding parameters in the P+T method. Typically PRS analysis involves constructing multiple PRSs across a range of the tuning parameters, followed by selection of the optimal PRS for prediction (i.e. the one that gives the strongest evidence for association). This optimization can inflate the probability of a type I error, if the multiple testing inherent in choosing the best PRS is not accounted for. Inflated type 1 error can be guarded against by using permutations to evaluate significance of the selected PRS; we refer to this approach as Opt-perm. Here, while the p-value would be corrected for multiple testing, the optimized PRS may still be over-fit and thus the corresponding R^2^ value, which measures the proportion of variation in the trait explained by the PRS, would be inflated. Moreover, it should be noted that permutation procedures can become quite computationally intensive. It has also been proposed to use external or internal validation to choose tuning parameters and avoid permutations. However, external validation datasets are often not available, especially for rarely-studied phenotypes^7^, and in smaller samples, splitting the data into training and validation sets can decrease power. As an alternative to the optimization approach, one could *a* priori choose a single tuning parameter setting (e.g. fixing the p-value threshold and LD pruning level) to construct a single PRS. This approach was used recently in two different investigations to test for association of one PRS with many phenotypes^8,9^. By not optimizing over a set of tuning parameters for each test of association, this strategy avoids further increasing the multiple testing and computation time. However, a sub-optimal PRS may be selected, leading to poor prediction and power to test for association of the PRS with the trait.

Here, we instead compute PRSs over a range of tuning parameter settings, perform principal component analysis (PCA) on the set of PRSs, and use only the first PRS-PC for association testing. The first PC captures the largest amount of variation in the computed PRSs and thus could have better discrimination of the phenotype we are testing. This strategy was recently implemented in a study of rare copy number variation and polygenic risk of schizophrenia^10^. This unsupervised approach incorporates all computed scores across a range of tuning parameters and, importantly, is ignorant of the outcome of interest and thus maintains correct type I error. Additionally, the PRS-PCA approach produces a score that is not overfit, which can be used to assess predictive performance of the PRS using measures such as R^2^ or area under the receiver operating characteristic curve (AUC).

Here, we assess the statistical properties of the proposed method in the context of P+T PRS analysis. We begin by constructing PRSs using the P+T approach across a range of p-value thresholds. We then compare the performance of the PRS-PCA approach with the Opt-perm approach and *a* priori selection of the p-value threshold tuning parameter. Using simulations and analysis of the Mayo Clinic Bipolar Disorder (BD) Biobank data, we show that the PRS-PCA approach maintains correct type I error and outperforms the other PRS strategies in most scenarios.

## Methods

### Polygenic risk scores

Let *G_ij_* denote the number of copies of the reference allele for the *j^th^* SNP for the *i^th^* individual, possibly estimated via imputation. Let 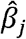 be the estimated effect for the *j_th_* SNP. The PRS for the *i^th^* individual is then 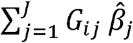 over a set *J* markers. Using GWAS summary statistics from a prior analysis of a trait of interest and LD structure estimated either from a reference panel or the target data, the set of *j* markers to include in the sum and their estimated effects are usually chosen using a P+T strategy^3^. Briefly, P+T “prunes” the genome to obtain approximately independent SNPs and only uses SNPs below a certain p-value threshold to estimate the PRS. This approach can be optimized over different p-value thresholds, clump sizes, and LD measures to determine approximate independence.

However, by searching for the best tuning parameter setting, the PRS can be overfit to the target data and the test of association of the PRS with a trait can have inflated type-I error. This can be corrected by using permutations to generate empirical p-values for association with the optimized PRS.

### PRS-PCA approach

Instead of using the target data to choose the best PRS, we propose using an unsupervised approach to construct a single PRS from the set of PRSs computed over a range of tuning parameter values (e.g. over a range of p-value thresholds). We first standardize each of the original K PRSs (corresponding to K different P+T settings) to have mean 0 and standard deviation 1, and construct a matrix [*PRS*_1_ *PRS*_2_..., *PRS_K_*] containing the K standardized PRSs. We then perform PCA on this matrix to obtain K independent PRS-PCs, which are weighted summations of the columns of the matrix. Just like a typical PRS, each PRS-PC is a weighted summation of the SNPs; however, the weights of the SNPs are different than in a standard PRS constructed using P+T. Specifically,

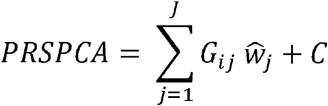

where 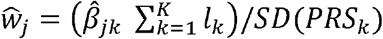, *l_k_* is the PCA loading for the *k^th^* PRS, 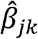 is the estimated effect of the *j^th^* SNP under setting *k* which for P+T is either 0 or the effect estimate from the source GWAS, and where *SD*(*PRS_k_*) and *C* account for standard deviation and mean, respectively, of the PRSs that are standardized before performing PCA. We keep only the first PRS-PC from the PCA which explains the greatest variation of the PRSs computed under different settings, and use it to test for association with the phenotype.

### Simulations

To estimate empirical type I error and power of the previous and newly proposed methods under different scenarios, we simulated data with or without genotype-phenotype associations. We generated genotypes by sampling without replacement from the Mayo Clinic Bipolar Disorder Biobank sample, followed by generating phenotypes conditional on (or independent of) the genotypes. The Mayo Clinic Bipolar Disorder Biobank collection, genotyping, and genetic data quality control has been described in previous publications^11,12^ and is summarized in the supplement.

We explored the performance of the methods using samples sizes of N = 500 and 1500 with a balanced case-control design. Using GCTA^13^, we simulated the liability of a trait with realistic effect sizes across the genome by randomly choosing the effect size of each SNP from a normal distribution with mean equal to log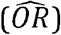 and standard deviation 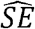 of the corresponding SNP in the summary statistics from the Psychiatric Genomics Consortium (PGC) Schizophrenia (SZ) GWAS^14^ which we previously showed was associated with psychosis during mania in bipolar disorder^11^. To avoid assigning “causal” effects to SNPs in LD, we first clumped the summary statistics using PLINKv1.90 (--clump-kb 250 -- clump-p=1 --clump-r2=0.1) to obtain 93 802 approximately independent SNPs. Additionally, we varied the level of polygenicity of the trait by choosing SNPs with absolute value of the log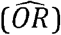 greater than 0.01 (high; 71694 SNPs), 0.07 (medium; 1493 SNPs), and 0.15 (low; 31 SNPs), respectively, in the PGC-SZ GWAS. We varied the heritability of the liability to be 0, 0.2, 0.4, 0.6, or 0.8. The final simulated liability was then dichotomized at the median to create a balance of cases and controls.

We used PRSice2^3^ to perform pruning and thresholding to compute PRSs in the simulated datasets using the PGC-SZ summary statistics. The simulated datasets were then analyzed using PRS-PCA as well as Opt-perm, and *a* priori selection of a p-value threshold (*p_T_* = 5*x*10^−8^, 0.05, or 1). To explore the effect of the number of p-value thresholds searched on the performance of PRS-PCA and optimization, we computed PRSs at either K = 5 (*p_T_* = 5*x*10^−8^, 10^−6^, 10^−4^, 0.01, 1), 11 (*p_T_* = 5*x*10^−8^, 10^−7^, 10^−6^, 10^−5^, 10^−4^, 0.001, 0.01, 0.05, 0.1, 0.5.1), or 106 (*p_T_* = 5*x*10^−8^, 10^−7^, 10^−6^, 10^−5^, 10^−4^, 0.001, 0.01, 0.02, 0.03, …,0.99, 1) different p-value thresholds. It should be noted that the default search implemented in PRSice2 searches multiple hundreds of p-value thresholds over an even grid from 5*x*10^−8^ to 1. We chose a smaller grid to reduce computational expense in our simulations. 15000 and 1000 replicates were used to estimate empirical type I error (heritability = 0) and power (heritability > 0), respectively, for each scenario.

### Application to Mayo Clinic Bipolar Biobank Data

To compare the performances of the PRS approaches, we used publicly available GWAS summary statistics to calculate PRSs for a variety of traits for subjects in the Mayo Bipolar Biobank dataset, including: SZ^14^, BD^15^, major depressive disorder (MDD)^16^, attention deficit and hyperactivity disorder (ADHD)^17^, anxiety disorders^18^, post-traumatic stress disorder (PTSD)^19^, obsessive compulsive disorder (OCD)^20^, anorexia nervosa (AN)^21^, insomnia^22^, and educational attainment (EA)^23^. We used PRSice2^3^ to compute the PRSs at 11 p-value thresholds (*p_T_* = 5*x*10^−8^, 10^−7^,10^−6^, 10^−5^, 10^−4^, 0.001, 0.01, 0.05,0.1, 0.5, or 1) with fixed pruning parameters (--clump-r2 0.1 and --clump-kb 250). We used the various PRS approaches to test for association of each PRS with the history of psychosis during mania in BD cases. We recently demonstrated that psychosis during mania is associated with polygenic risk of schizophrenia^11^. No large GWAS exists for this phenotype, thus, PRS approaches can be quite useful here to elucidate potential differences in genetic background between bipolar cases with and without psychosis, and the genetic overlap of this phenotype with other psychiatric traits in addition to SZ. We used logistic regression to test for association of each PRS with psychosis status after controlling for the first four principal components of the genotype data to adjust for population stratification. 100,000 permutations were used to calculated p-values for the Opt-perm method. We estimated the proportion of variation of the binary phenotype explained by each PRS using Nagelkerke’s R^2^. For the Opt-perm approach, we followed the standard approach of reporting the Nagelkerke’s R^2^ estimate for best p-value threshold, which is a biased overestimate of the true R^2^.

## Results

### Type I error

Table 1 shows the empirical type I error for each method, corresponding to setting the heritability of the liability equal to zero. A total of 15000 simulations were performed for each scenario (row) in Table 1. The PRSs with *a* priori p-value threshold and the PRS-PCA approach maintain correct type I error in all scenarios. As expected, optimization of p-value threshold without correction for multiple testing results in inflated type I error, which worsens as the number of thresholds searched increases. Permutations correct the type I error.

**Table 1.** Empirical Type I error for each method with sample size N and number of parameters searched K. PCA = first PC of search, Opt = Select best parameter, Opt-perm = Permutation of Opt

| N | K | p = 5e-8 | p = 0.05 | p = 1 | PCA | Opt | Opt-perm |
| --- | --- | --- | --- | --- | --- | --- | --- |
| 500 | 11 | 0.049 | 0.048 | 0.049 | 0.049 | 0.187 | 0.050 |
| 500 | 106 | 0.047 | 0.052 | 0.046 | 0.045 | 0.208 | 0.048 |
| 1500 | 11 | 0.050 | 0.049 | 0.047 | 0.050 | 0.189 | 0.050 |
| 1500 | 106 | 0.051 | 0.049 | 0.051 | 0.053 | 0.204 | 0.051 |

### Investigation of the PRS-PCA approach

Figure 1 shows a comparison of SNP weights between the PRS-PCA score computed using either K = 11 or 106 thresholds, and the PRSs with specific p-value thresholds (*p_T_* =5e-8, 0.05, or 1). Using p-value thresholds less than 1 sets some of the SNP weights equal to zero and thus excludes those SNPs from the PRS. Similar to choosing a p-value threshold of 1, the PRS-PCA approach assigns weight to all SNPs after pruning. However, SNPs with larger p-values are down-weighted in the PRS-PCA approach. When K is large (e.g. K = 106), SNPs with larger p-values are down-weighted less heavily and the PRS-PCA weights are almost proportional to weights for the PRS with p-value threshold of 1.

**Figure 1.**
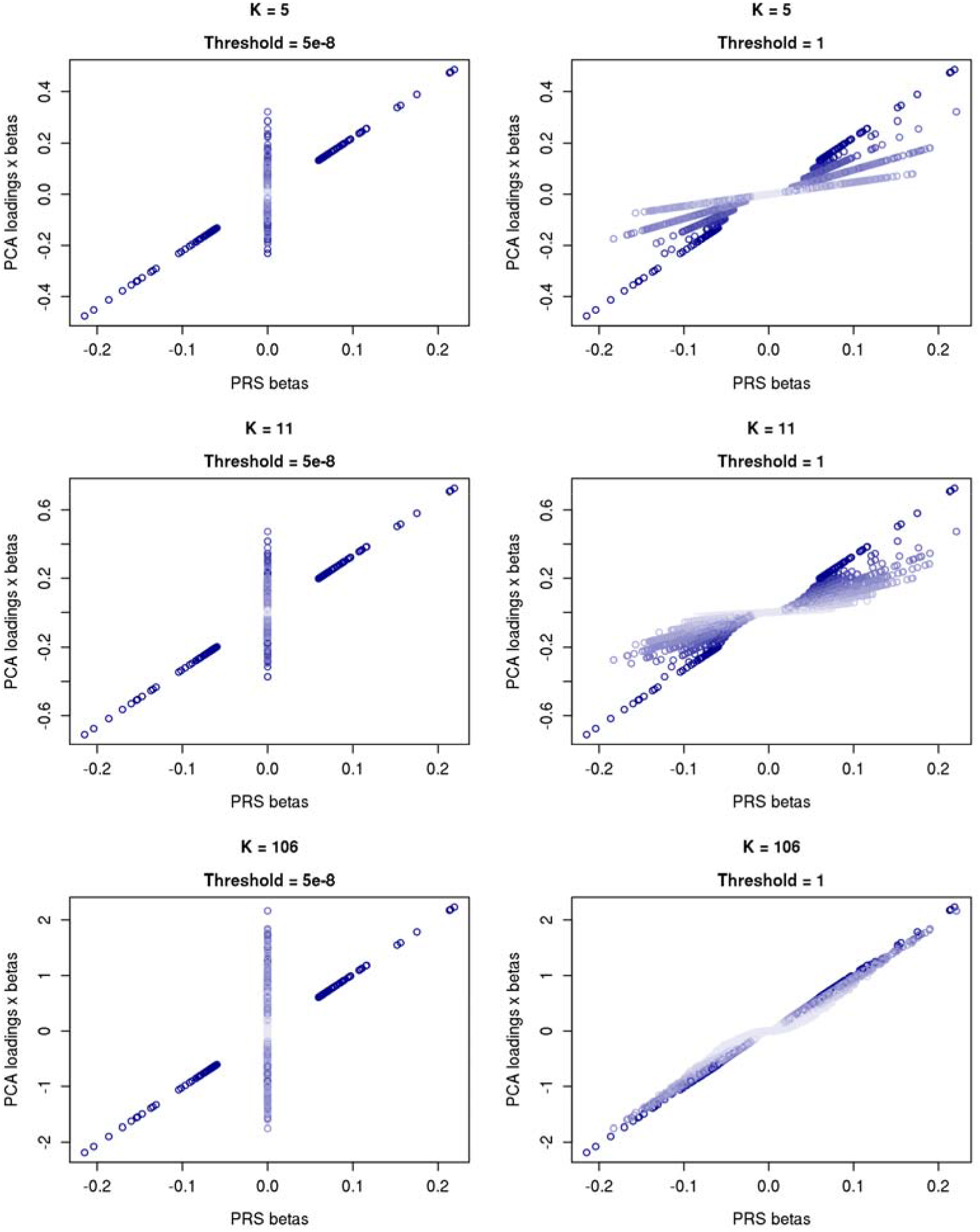
Comparison of SNP weights for SZ-PRS between PRS-PCA (y-axis) and PRS with p-value threshold of genome-wide significant (left) or 1 (right). Points are shaded based on p-value from SCZ GWAS (Dark blue: p < 5e-8; White: p = 1)

### Power

We assessed the power to detect association using the various PRS approaches for a trait with high, medium, or low polygenicity using 1000 simulations for a given sample size and heritability. Empirical power for these methods is shown in Figure 1. All methods had more power using a sample size of 1500, but the relative performance of the methods remained unchanged between sample sizes. The PRS-PCA approach had competitive power in all scenarios with K = 5 or 11 and nearly achieved the same power as the PRS constructed with fixed threshold matching the simulation setting (*p_T_* = 5e-8 and 1 for low and high polygenicity, respectively). PRS-PCA performed substantially worse when using many p-value thresholds (K=106) because SNP weights mimic the PRS constructed with all SNPs. PRS=PCA with this choice of K performs only marginally better than PRS including all SNPs. The Opt-Perm approach performed similarly regardless of the number of threshold searched, and had similar or less power than PRS-PCA approaches with K = 5 or 11.

### Illustration of Approach: Application to Mayo Clinic Bipolar Biobank Data

Figure 3 displays the proportion of PRS variation explained by each PRS-PC as well as the PCA loadings of the fixed-threshold PRSs in the first PRS-PC, for both K = 5 and 11. For the psychiatric traits considered, the first PRS-PC explained between 40% and 74% of the variation in PRSs computed at different p-value thresholds regardless of K. Table 2 shows the results for the PRS analyses. The PRS-PCA method with K = 11 showed that the PRSs for EA, BD, and SZ were higher in cases with psychosis than those without. With K = 5, the best threshold was left out of the search for the PRS-PCA approach and thus it lost power. The Opt-Perm method (with K = 11) provided weaker evidence of association with the BD-PRS, but stronger evidence of a PTSD PRS association than the PRS-PCA method.

**Figure 2.**
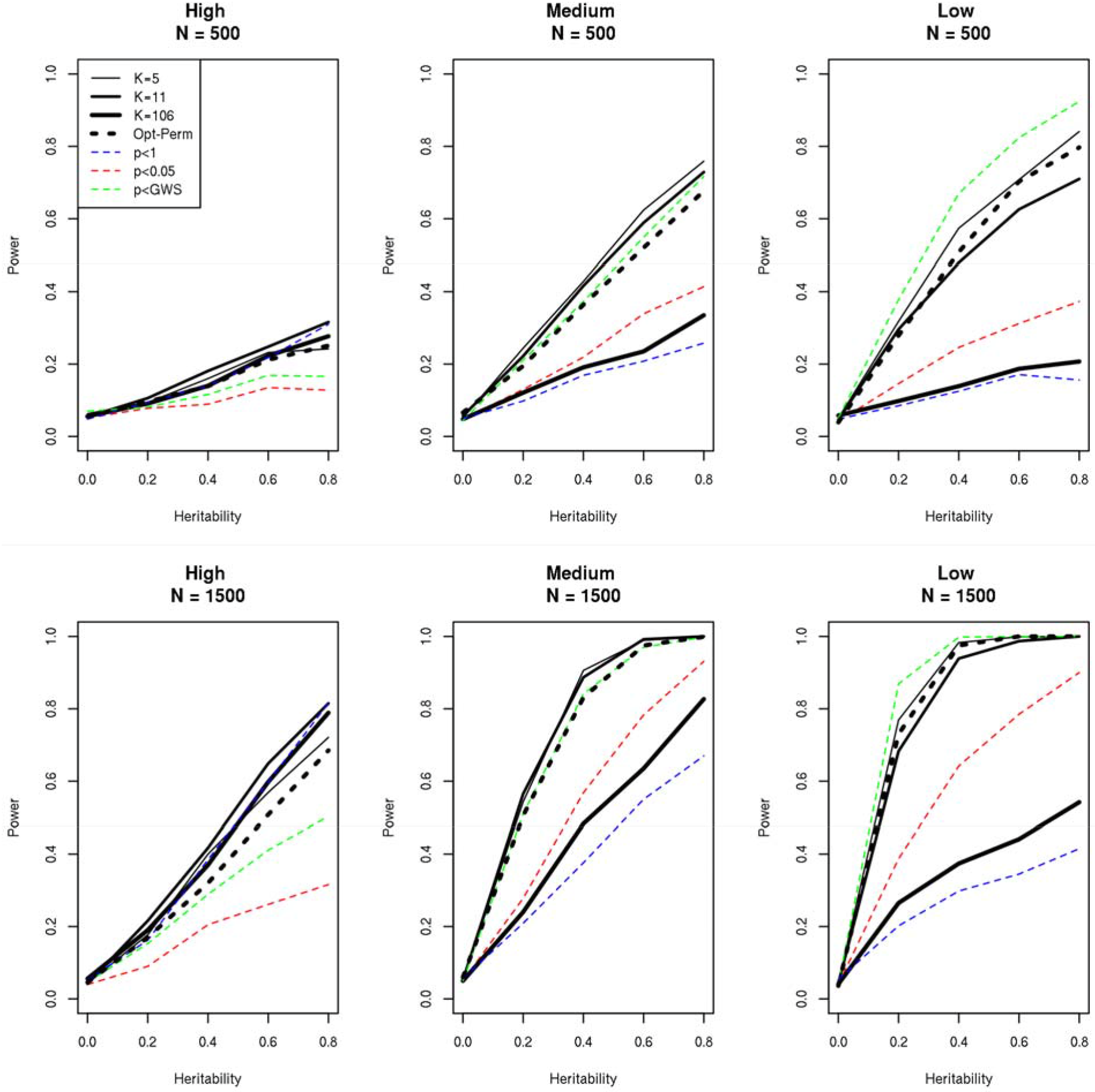
Empirical power of each method given a trait with high (left; |log(OR)| > 0.01), medium (center; |log(OR)| > 0.07), or low (right; |log(OR)| > 0.15) polygenicity with sample size of N = 500 (top) or 1500 (bottom). K = PRS-PCA using K PRSs, GWS = genome-wide significant p-value threshold (5e-8).

**Figure 3.**
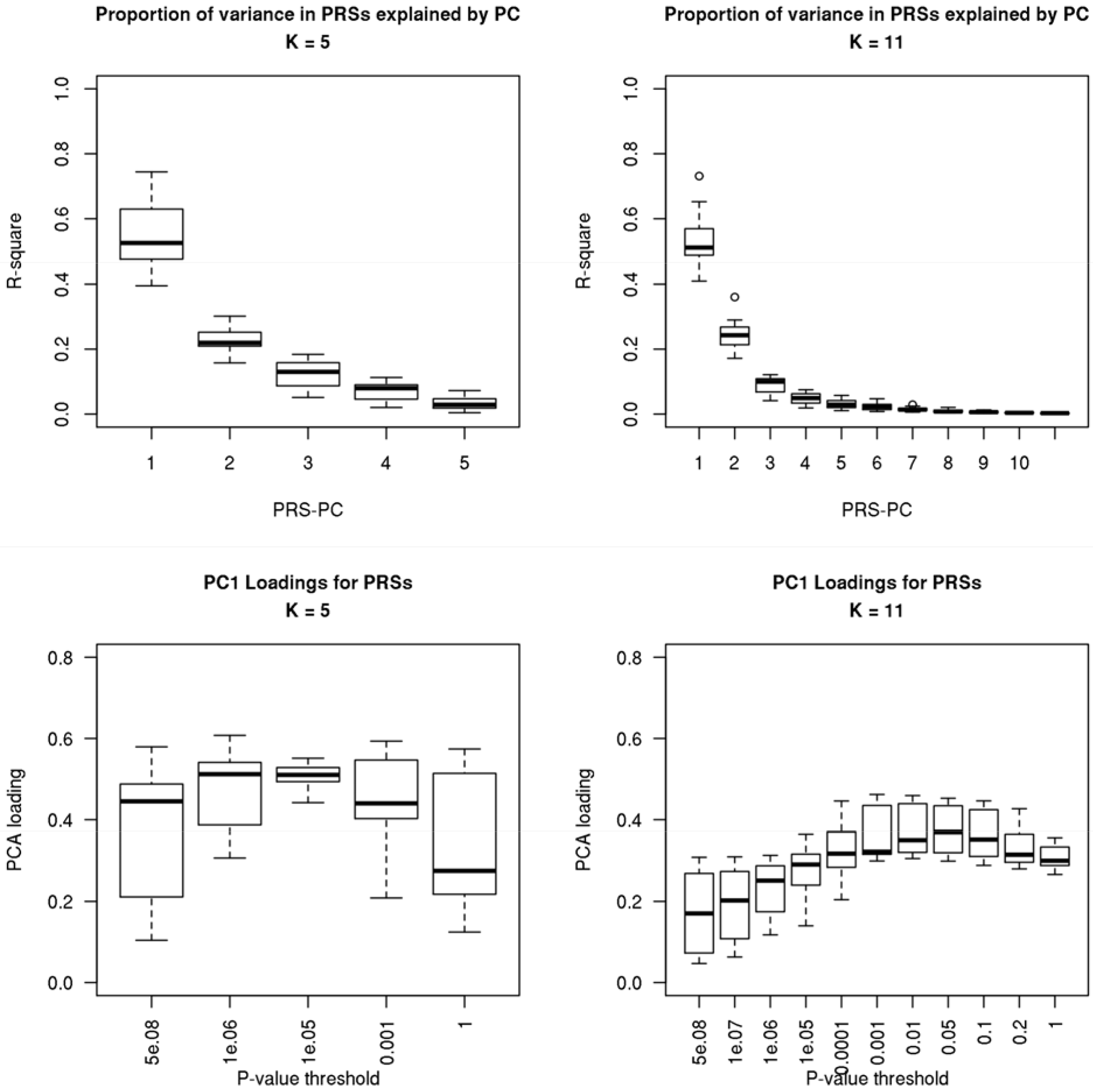
Boxplots summarizing the proportion of variation in PRSs explained by each PC (top) and the loadings in the first PC (bottom) of PRSs at each threshold, for the PRSs analyzed in Table 2.

**Table 2.** Comparison of PRS approaches testing for association of each PRS with presence of psychosis among cases of bipolar disorder (N = 645). The traits are sorted by the PCA approach (K = 11) p-value. Prediction performance measured by Nagelkerke’s R^2^. *Nagelkerke’s R^2^ for the Opt-Perm approach is estimated from the best performing p-value threshold searched.

| PRS | PC1 Prop. Variance explained | PCA (K = 5) p-value ($R^2$) | PCA (K = 11) p-value ($R^2$) | Opt-Perm p-value ($R^2$)* |
| --- | --- | --- | --- | --- |
| EA | 73% | 0.001 (2.3%) | 0.001 (2.2%) | 0.006 (2.2%) |
| BD | 49% | 0.013 (1.1%) | 0.020 (1.0%) | 0.132 (1.0%) |
| SZ | 65% | 0.141 (0.3%) | 0.045 (0.7%) | 0.013 (1.8%) |
| PTSD | 52% | 0.623 (0%) | 0.115 (0.4%) | 0.041 (1.4%) |
| Anxiety | 46% | 0.459 (0%) | 0.270 (0.1%) | 0.829 (0.2%) |
| AN | 50% | 0.732 (0%) | 0.551 (0%) | 0.075 (1.1%) |
| OCD | 52% | 0.674 (0%) | 0.584 (0%) | 0.888 (0.1%) |
| ADHD | 49% | 0.806 (0%) | 0.671 (0%) | 0.664 (0.3%) |
| MDD | 50% | 0.890 (0%) | 0.718 (0%) | 0.438 (0.5%) |
| Insomnia | 48% | 0.618 (0%) | 0.884 (0%) | 0.350 (0.6%) |

## Discussion

In this paper, we proposed a method of PRS analysis that uses PCA to concentrate the maximum variation in a set of PRSs in a single PC, and then tests for association of the phenotype with only the first PC. This method avoids optimizing the parameters to construct the PRS, which inflates the probability of a type I error if unaccounted for, and is computationally faster than using permutations to correct for the inflation. Through simulations, we showed that the PRS-PCA approach with K = 5 or 11 can be as or more powerful than the Opt-Perm approach (with p-value computed using permutations). When a large grid search of p-value thresholds was used, the PRS-PCA approach mimicked the weights of the PRS including all SNPs and thus lost power in less polygenic models. In the BD data application, the PRS-PCA and Opt-Perm approaches obtained similar results. In addition to being computationally faster than the Opt-Perm approach, because PRS-PCA tests a single PRS rather than selecting the most predictive PRS in a particular dataset, the PRS-PCA approach produces an unbiased estimate of PRS performance (e.g. area under the curve or proportion of variation explained). While the performance of the PRS-PCA approach depends on the tuning parameter grid search used, our results suggest an even search over the negative log-ten p-value space performs well. Our search with K = 11 is a very typical choice in the field, and similar searches have been used previously^10^. This choice performed well in the simulations and, unlike the PRS-PCA approach with only K = 5, was able to reproduce the finding of Markota *et al*.^11^.

Both the PRS-PCA (K = 11) and the Opt-Perm approaches reproduced our previous finding that the PRS for SZ is higher in cases with a history of manic psychosis (N = 336) than those without a history of psychosis (N = 309)^11^. Both methods also showed cases with manic psychosis had higher genetic load for educational attainment. While psychosis in the context of bipolar disorder has been less studied, prior studies have shown small positive genetic correlation of educational attainment with SZ and the PRS for EA has been found to be higher in cases of SZ^24^. Finally, only the PRS-PCA approach found evidence that the PRS for BD was higher in cases with manic psychosis. This could reflect that a higher genetic load for BD can cause more severe symptoms of BD. This could also occur if the cases in the PGC study of BD had higher prevalence of psychosis and thus the training data better reflects cases with psychosis rather than without.

The PRS-PCA approach was designed to control type I error while maintaining good power. This approach is most suited to hypothesis testing with many PRSs because it prevents overfitting each PRS to the outcome and does not require choosing one p-value threshold for all PRSs^8,9,25^, which can reduce power. In this paper, we explored how the PRS-PCA approach can improve PRS analyses that implement P+T. Future investigation is needed to test if the same PCA approach can be used to avoid optimizing over different sets of tuning parameters with non-P+T PRS approaches^4–6^, such as lassosum^4^, LDpred^5^, or PRS-CS^6^. Furthermore, there is no uniformly most powerful method to construct PRSs and PRSs constructed under different methods could easily be combined using the PCA approach. This will be investigated in the future.

In this paper, we propose a new powerful method of testing for association of PRSs with a phenotype, which avoids the multiple testing inherent in the popular optimization approach. In studies that aim to test for association of PRSs with more than one phenotype such as a PRS PheWAS^8^, the PRS-PCA approach would substantially reduce the multiple testing that would occur with the optimization approach. With the growing use of PRSs, the PRS-PCA approach gives researchers an unbiased and powerful approach to dissect polygenic risk of phenotypes.

## Acknowledgments

This work was supported by the Marriott Foundation and the Mayo Clinic Center for Individualized Medicine.

## Conflicts of Interest

The authors have no conflicts of interest to report.

